# Experimental evolution of multicellularity via cuboidal cell packing in fission yeast

**DOI:** 10.1101/2023.11.03.565576

**Authors:** Rozenn M. Pineau, Penelope C. Kahn, Dung T. Lac, Mia Denning, Whitney Wong, William C. Ratcliff, G. Ozan Bozdag

## Abstract

The evolution of multicellularity represents a major transition in life’s history, enabling the rise of complex organisms. Multicellular groups can evolve through multiple developmental modes, but a common step is the formation of permanent cell-cell attachments after division. The characteristics of the multicellular morphology which emerges has profound consequences for the subsequent evolution of a nascent multicellular lineage, but little prior work has examined these dynamics directly. Here we examine a widespread yet understudied emergent multicellular morphology: cuboidal packing. Extinct and extant multicellular organisms across the tree of life have evolved to form groups in which spherical cells divide but remain attached, forming approximately cubic subunits. To experimentally investigate the evolution of cuboidal cell packing, we used settling selection to favor the evolution of simple multicellularity in unicellular, spherical *Schizosaccharomyces pombe* yeast. Multicellular clusters with cuboidal organization rapidly evolved, displacing the unicellular ancestor. These clusters displayed key hallmarks of an evolutionary transition in individuality: groups possess an emergent life cycle driven by physical fracture, group size is heritable, and they respond to group-level selection via multicellular adaptation. In 2/5 lineages, group formation was driven by mutations in the *ACE2* gene, preventing daughter cell separation after division. Remarkably, *ACE2* mutations also underlie the transition to multicellularity in *Saccharomyces cerevisiae* and *C. galabrata*, lineages last shared a common ancestor *>*300 million years ago. Our results provide insight into the evolution of cuboidal cell packing, an understudied multicellular morphology, and highlights the deeply convergent potential for a transition to multicellular individuality within fungi.

## 1 INTRODUCTION

Multicellularity evolved at least 50 times independently in different lineages, underscoring the diverse evolutionary paths that may lead to a multicellular life history ^1,2^. Over the last several decades, a broad range of work has highlighted just how influential the traits of the unicellular ancestor are during the transition to multicellularity (^3,4^). For example, the evolution of cellular specialization is thought to arise via co-option of temporally-varying cellular phenotypes, putting the unicellular toolkit to novel multicellular use (^5,6,7^). Similarly, the morphology of multicellular groups is heavily dependent on the traits of the unicellular ancestor. How cells divide and attach to one another has fundamental consequences for the biophysical properties and evolutionary dynamics of these lineages ^8,9^.

In this paper, we focus on early steps in evolution of clonally-developing multicellular morphologies, which result from the formation of permanent bonds between mother and daughter cells. If a cell can bud multiple times, remaining attached to each offspring, like the yeast *Saccharomyces cerevisiae*, then mother-daughter attachment results in branched groups with a tree-like morphology (e.g., snowflake yeast, ^10^, marine yeast ^11^, algae ^12^). Alternatively, if the cells divide via binary fission, and use the same division plane repeatedly, then permanent mother-daughter bonds result in linear filaments (*e*.*g*., *Nostoc*). If cells with binary fission instead have alternating division planes, then 3D structures with repeating cubic subunits of cells (composed of tetrads or octads) ^13^ can result-a morphology that has been termed “cuboidal packing” ^14^ (see Figure A.3 for a schematic of cuboidal packing).

Cuboidal packing has evolved repeatedly across the tree of life (Figure 1), in both contemporary and ancient organisms. For example, one of the most enigmatic macroscopic multicellular fossils from the Doushantuo Formation in southern China (570 MYA) forms large groups with cuboidally-packed units (Figure 1 A, panel 1). This species, which has been hypothesized to be an algae ^14^, grows similarly to modern day chlorosarcinacean green algae, like the lichen-forming green algae *Diplosphaera chodatii* (Figure 1 A, panel 2). Cuboidal packing also appears within both the archaea and bacteria, for instance, in anaerobic methanogens and in the bacterial genus *Sarcina spp*. (*Methanosarcina spp*., Figure 1 A, panel 3, *Sarcina spp*. Figure 1 A, panel 4). While cuboidal packing is a relatively common route for group formation, no prior work has directly examined how it arises *de novo*, and whether it can support an evolutionary transition in individuality in which groups of cells reproduce, possess heritable variation in emergent multicellular traits, and respond to group-level selection with multicellular adaptation.

**FIGURE 1.**
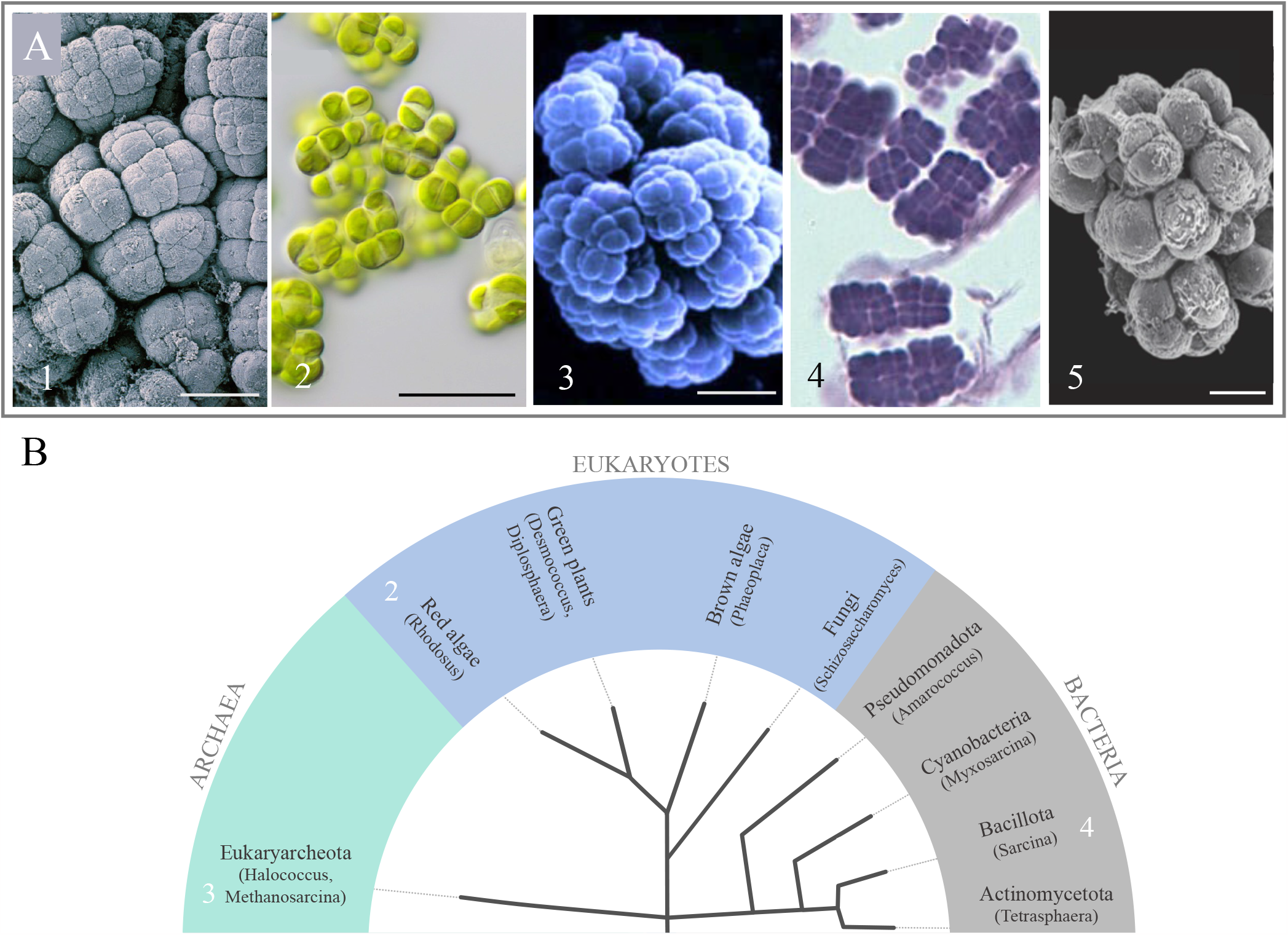
Cuboidal patterning is found in all major branches of the tree of life. (A) Cuboidal packing has been observed in (1,5) fossils (Doushantuo algal thalli ^14^, *Archeophycus yunnanensis* ^15^), and (2) extant eukaryotes (*Diplosphaera chodatii*) ^16^, in (3) archaea (*Methanosarcina spp*.) ^17^ and (4) bacteria (*Sarcina spp*.) ^18^. The scale bar is 20 *μ*. (B) Cladogram depicting the phylogenetic relationship of the examples above. See Supplementary Table 1 for details on these taxa.

Here we investigate the evolution of cuboidal cell packing, a widespread yet understudied multicellular morphology, using spherical fission yeast as a model system. We subject these cells to settling selection, which favors larger group size, and observe the emergence of multicellular clusters with cuboidal organization within 20 rounds of selection (it is subsequently maintained until the end of the experiment, at 334 rounds of selection). We characterize the life cycle, heritability, and fitness of these clusters, and show that they gain the capacity for multicellular adaptation, satisfying the key criteria for an evolutionary transition in individuality. We also identify the genetic basis of cluster formation, and find that mutations in the *ACE2* gene, which prevents cell separation after division, are responsible for this phenotype in two independent lineages. Our findings illustrate the simplicity and convergence of the genetic and biophysical factors that enable the origin of multicellular groups and life cycles and shed light on the evolution of cuboidal cell packing across the tree of life.

## 2 METHODS

### 2.1 Evolution experiment

We applied daily settling selection to examine the evolution of nascent multicellularity in the unicellular fission yeast, *Schizosaccharomyces pombe*. We started the selection experiment with an isogenic clonal ancestor of a spherical single-celled yeast (sph2-3 ade3-58 h90^19^) by growing it as five independent replicate populations in 5 mL YES media (0.5 % w/v yeast extract, 3 % w/v glucose, 225 mg/L adenine, histidine, leucine, uracil, and lysine). After every 24 hours of growth (30 °C, 220 rpm), we transferred 1 mL of the culture into a 1.5 mL Eppendorf tube. Following 5 minutes of settling through gravity, we discarded the top portion of the culture and transferred the bottom 25 *μ*L into a fresh 5 mL YES culture to start the next round of growth and settling selection. We applied a 5-minute settling selection for the first 190 days, followed by a more robust selection regime of 1-minute settling selection until the final time point of 334 days. We made 30 % glycerol stocks of each replicate population at approximately 15-day intervals and stored them at −80 °C for further examination.

### 2.2 Measuring cluster biomass and area

To measure the increase in multicellular size, we picked single colony isolates from all five populations at t60, t125, and t344 (t=days). We then grew these isolates for three days with growth and settling selection, replicating the same conditions as our experimental evolution protocol. To measure the area of these evolved clusters, we diluted them 4000 times and pipetted these 15 samples into 24-well plates and imaged the whole well with a Nikon Eclipse Ti microscope at 40x magnification. Using custom-generated MatLab scripts, we measured the area of each cluster in microns, obtaining at least 3000 data points per sample. Finally, to take glamour shots representing the ancestral and evolved *S. pombe* cells and clusters, we used a Nikon AR1 confocal microscope.

### 2.3 Time-lapse microscopy

To determine the growth and reproduction of evolved *S. pombe* clusters, we used time-lapse microscopy. Specifically, we first grew cultures overnight in 10 mL YES media. Next, we diluted them 10^3^ and 10^4^ times in 2x YES media combined with soft agar at a final concentration of 0.035%. We then pipetted 100 *μ*L of a mixture of soft agar and clusters into 8-well chamber slides. We took images of the growing culture every 20 minutes for 24 hours at a 200X magnification and across five different Z planes using a Nikon Eclipse Ti microscope. Finally, we processed these images using ImageJ.

### 2.4 Growth rate measurements

To examine the daily growth rate of evolved clusters, we revived single cluster isolates from all five frozen populations at t125 by growing them to equilibrium for three days under their original growth conditions. We started recording the growth rate (t0 time point) after one round of settling selection. We then sampled the culture after 3, 6, 12 and 24 hours of growth. We estimated population size (number of clusters) and cluster size distribution for each time point in 24-well plates by imaging the whole well with a Nikon Eclipse Ti microscope at 40x magnification.

### 2.5 Effect of selection assay

To assess whether heritable differences in cluster size led to a predictable difference in survival component of fitness (Figure 3 E), we compared the survival advantage of clusters at t334 against clusters at t60, the former being, on average, 20% larger in size. First, we selected single clonal isolates from each time point. Next, to be able to distinguish the frequency of the two genotypes using a replica plating assay, we genetically transformed the t60 isolate with a hygromycin resistance cassette (see details below). To start the experiment, we first inoculated each genotype separately in YES media overnight and mixed them in equal volumes with four technical replicates. We then plated these mixed cultures (with 4000- and 8000-fold dilutions) both before and after subjecting them to 5-mins of settling selection. After 3 days of growth, we replica-plated colonies from the original plates to plates containing hygromycin and let them grow for 2 days. Finally, we counted CFUs on YES and YES+Hyg plates for cultures plated before and after the settling selection, giving us the fraction of the two genotypes and the rate of increased survival due to larger size at t334. The selection rate was calculated as follow: *selection rate* = *t*334_*tf*_ /*t*334*t*_*t*0_ − *t*60_*tf*_ /*t*60_*t*0_.

### 2.6 Strain construction

To compete the t60 and t334 strains (Figure 3), we substituted the *URA4* gene with a hygromycin resistance cassette in a t60 strain isolate. Specifically, we inserted the hygromycin B resistance cassette at the *URA4* site *ura4*Li*::HYGMX//URA4* using the Lithium Acetate/Dimethyl Sulfoxide Procedure outlined by Murray et al. ^20^ and selected for mutants capable of growing on YES - hygromycin plates (see primers in Table 1). To obtain homozygous diploids of *ura4*Li, we induced sporulation by growing the population in YEPG for two days, followed by tetrad dissection and allowing haploid mutants to self-fertilize to form homozygous diploid mutants.

**TABLE 1.**
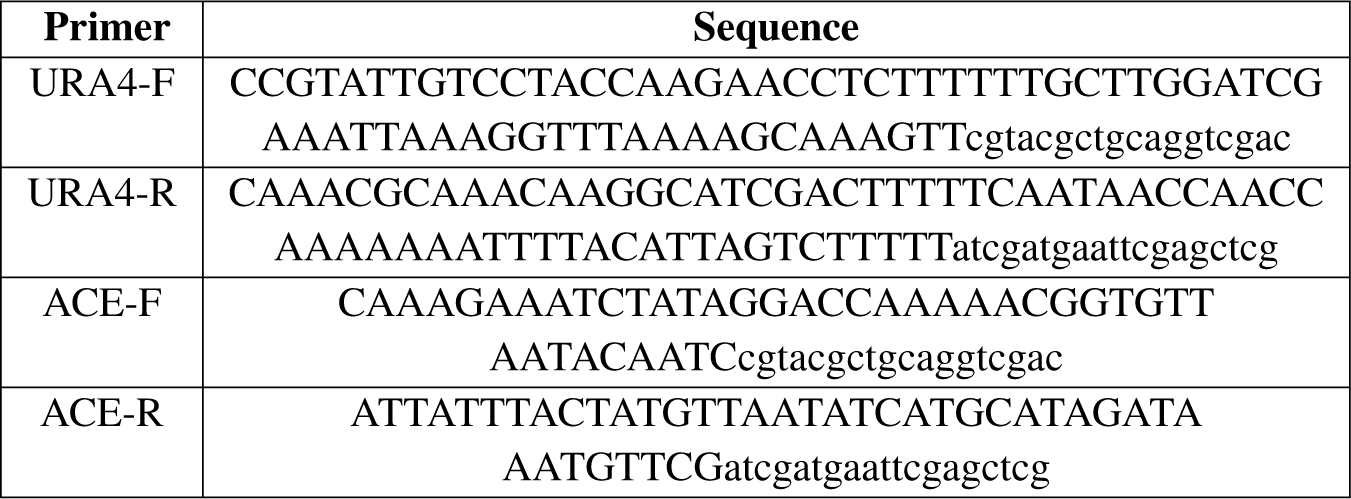
Primers sequences for *URA4* (see Figure 3) and *ACE2* (see Figure 4) gene knockouts with insertion of resistance to hygromycin. Uppercase letter corresponds to the gene-matching primer sequence, and the lower case letters to the plasmidmatching sequence.

To examine the phenotypic consequences of the loss of function in the *ACE2* gene (Figure 4), we deleted the *ACE2* open reading frame with a hygromycin resistance cassette. Using the same transformation protocol mentioned above, we inserted the hygromycin resistance cassette at the *ACE2* locus *ace2*Li*::HYGMX//ACE2*, and selected transformants growing on YES - hygromycin plates (see Table 1 for primer sequences). Finally, to obtain homozygous diploids (ace2Li::ace2Li), we induced sporulation by growing the population in YEPG for two days, followed by tetrad dissection and allowing haploid mutants to self-fertilize to form homozygous diploid mutants.

### 2.7 gDNA extraction and bioinformatics analysis

To identify candidate mutations for spherical cell shape and cluster formation, we sequenced clonal isolates of the unicellular ancestor and evolved clusters from five independent populations at t334. We used VWR’s Life Science Yeast Genomic DNA Purification Kit to extract genomic DNA. We then mailed the genomic DNA samples to the Microbial Genome Sequencing Center (SeqCenter, https://www.seqcenter.com/) for library preparation and Illumina sequencing.

We analyzed 150-bp paired-end Illumina reads on a Linux platform. First, we used FASTP (v.0.21.0^21^) to trim the first 15 and the last cycles of the sequencing run and filter out reads lower than a PHRED score of 30. We aligned reads to the most recent version of the *S. pombe* reference genome (https://www.pombase.org/data/genome_sequence_and_features/genome_sequence/) using BWA-MEM (v.0.7.17^22^). Next, to sort, index, and convert SAM files to BAM files and to mark the duplicates, we used PICARD tools (v.2.24.0, http://broadinstitute.github.io/picard/), SAMTOOLS (v.1.7^23^), and BAMTOOLS (v.2.3.0^24^). We then used GATK’s ValidateSamFiles and HaplotypeCaller tools (v.4.1.9.0) to validate BAM files and call variants ^25^. Next, we used VCFtools (v.0.1.17^26^) to filter out variants with a PHRED score lower than 30, a minimal depth below 12, and an allele frequency lower than 0.1. We then identified de novo variants unique to the evolved clusters using BCFTOOLS isec (v1.16). Finally, for a last round of filtering, we visually inspected VCF files on Integrative Genomics Viewer (IGV, v.2.8.13^27^) and removed variants at highly complex regions with alignment issues.

We identified candidate variants for the spherical cell shape by searching through the literature for known effects of the mutations detected in the sequenced ancestral strain (see Supplementary Table for a complete list of the mutations).

## 3 RESULTS

Multicellular organisms with cuboidal packing typically have spherical cells (see Figure 1a). This may facilitate tetrad and octad formation by lowering geometric constraints on cellular packing. We thus started our experiment with a spherical mutant clonal isolate of the fission yeast *Schizosaccharomyces pombe*. This round mutant phenotype was obtained by screening for spherical mutants after UV radiation by Sipiczki et al. ^19^. We sequenced its genome and we identified GEF1p as a potential candidate mutation for sphericity. GEF1p activates a pathway for apical growth and to mediate cytokinesis that involves *Cdc42*, and mutations inactivating *Cdc42* produce round spherical *S*.*pombe* cells (^28^ see supplementary table 1 for details on the mutations).

We evolved five replicate populations of initially clonal spherical *S. pombe* with daily settling selection-an experimental regimen shown to favor group formation in a range of organisms (snowflake yeast ^29^, *Sphaeroforma arctica* ^30^, *C. reinhardtii* ^31^, *Kluyveromyces lactis* ^32^), for a total of 334 transfers. In brief, we grew 5 mL of culture for 24 hours and transferred a randomly subsampled 1.5 mL into a microcentrifuge tube. After a 5-minute gravity settling selection, we discarded the top layer and transferred only the bottom 2.5% into a new 5 mL culture for the subsequent growth and settling phase(Figure 2 A).

**FIGURE 2.**
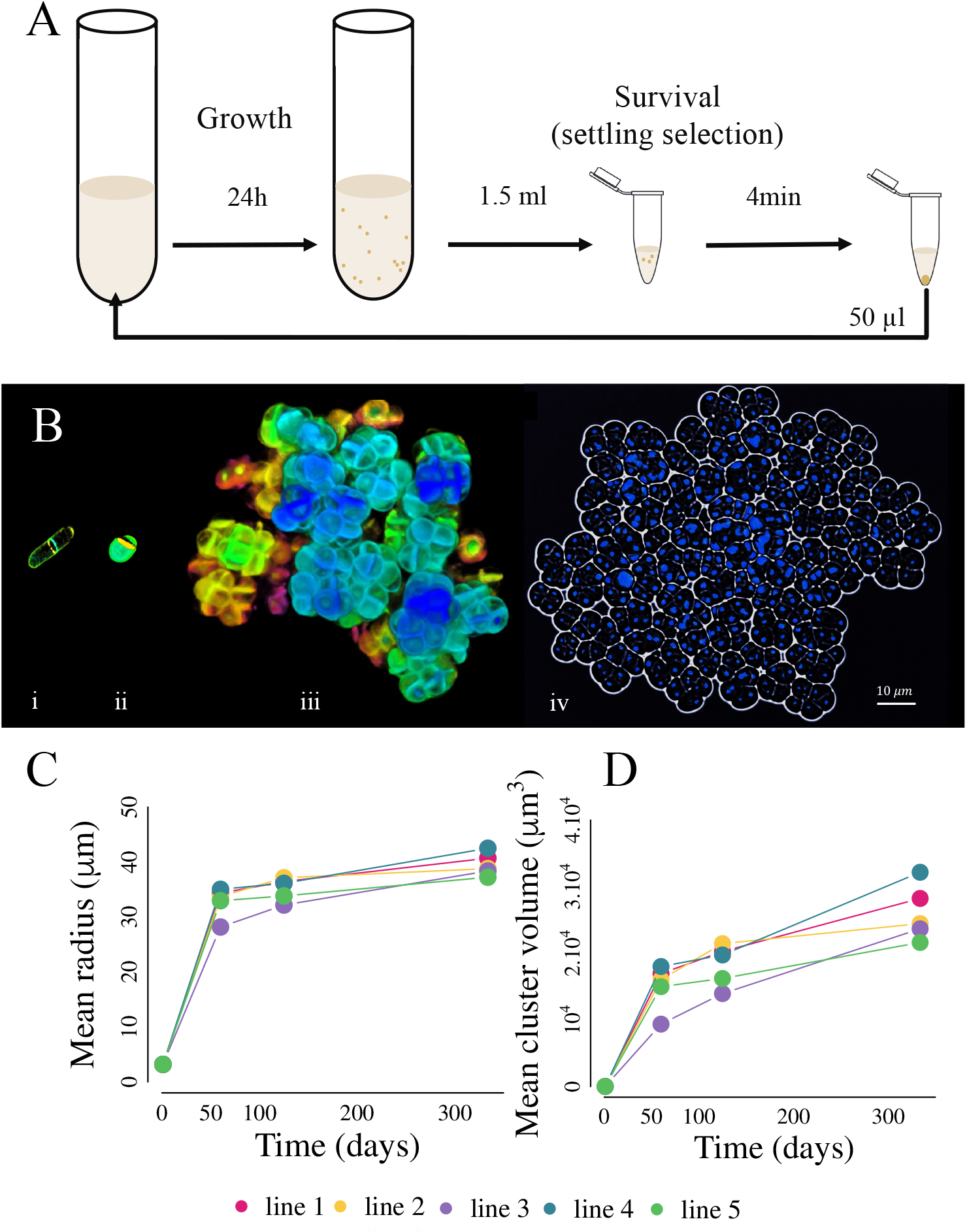
Emergence of multicellular groups in spherical fission yeast. (A) Culture cycle. Each day, *S. pombe* was grown for 24 hours before being subject to settling selection for larger size. (B) Cuboidally-packed multicellular groups evolved within 20 days. The image shows the wild-type ancestor (i), the spherical single-celled mutant (ii), and day-20 multicellular *S. pombe* in confocal microscopy (iii) and fluorescence microscopy (iv) with DAPI-stained nuclei. Colors in (iii) represent depth in z-position. See Figure A.1 for images of whole populations for the five independently evolving lines. (C) The size of the clusters increases by a factor of 10 in the first 60 days (∼140 generations), followed by a slower increase (1.2-fold in the last 274 days, ∼790 generations total).

This approach simultaneously favors three traits essential for the evolution of multicellularity: first, settling selection favors group formation, as groups settle faster than single cells. Only the fastest settling 2.5% of the sub-population’s biomass is transferred, thus competition is an open-ended arms race (*i*.*e*., there is no size at which survival is guaranteed). Second, by discarding 80% of the population prior to settling selection, we maintain strong selection on there being a multicellular life cycle in which groups reproduce. That is, a lineage which simply grows indefinitely without reproducing will eventually be discarded, and is an evolutionary dead end. Finally, the 24 h of growth maintains strong selection for rapid growth. These traits may trade-off with one another, and collectively ensure that we have created conditions under which groups that serve as multicellular Darwinian entities may arise.

Within 20 days of evolution, we observed the emergence of groups with a cuboidal packing morphology (Figure 2 B), and by day 60 (∼140 generations), the multicellular clusters had outcompeted their unicellular ancestor. We observed a 10-fold increase in cluster size within the first 60 days of the experiment, coinciding with the initial formation of multicellular groups. This was followed by a more gradual, 1.2-fold increase in group size over the remainder of the experiment from day 60 to day 334 (∼790 generations total, Figure 2 C&D). Differences in size between the four time points (day 0, 60, 125, and 334) were statistically significant based on a two-way ANOVA (_2,72789_ = 3430, *<* 2.10^−16^ for the effect of size, _2,72789_ = 46.92, *<* 7.10^−12^ for the effect of population) with post-hoc Tukey’s HSD tests (all pairwise comparisons significant at *<* 0.01).

Next, we examined the capacity for these groups to serve as Darwinian individuals ^33,34^. This occurs when groups reproduce, forming new groups, and there is heritable variation in group-level traits that affects fitness. When these conditions are met, multicellular Darwinian individuals can gain multicellular adaptations via natural selection, which can ultimately lead to the evolution of functionally-integrated multicellular organisms ^35^.

To explore the life cycle of multicellular *S*.*pombe*, which is required for groups to beget new groups, we imaged clusters from line 3 at t334 over a 24-hour period in soft agar. Clusters grew until a multicellular propagule separated from the main group via spontaneous fragmentation (Figure 3 A), forming new groups from these propagules. Over 24 h of growth, their size distribution remained remarkably stable (mean of ≈ 15 *μ*m over the 24 h growth cycle, Figure 3 B), while the number of clusters increased super-linearly (the mean growth rate was 0.13 clusters per hour across all five lines, Figure 3 C). Multicellular *S. pombe* thus have an emergent multicellular life cycle generated by growth and reproduction, the latter driven by spontaneous fracture.

**FIGURE 3.**
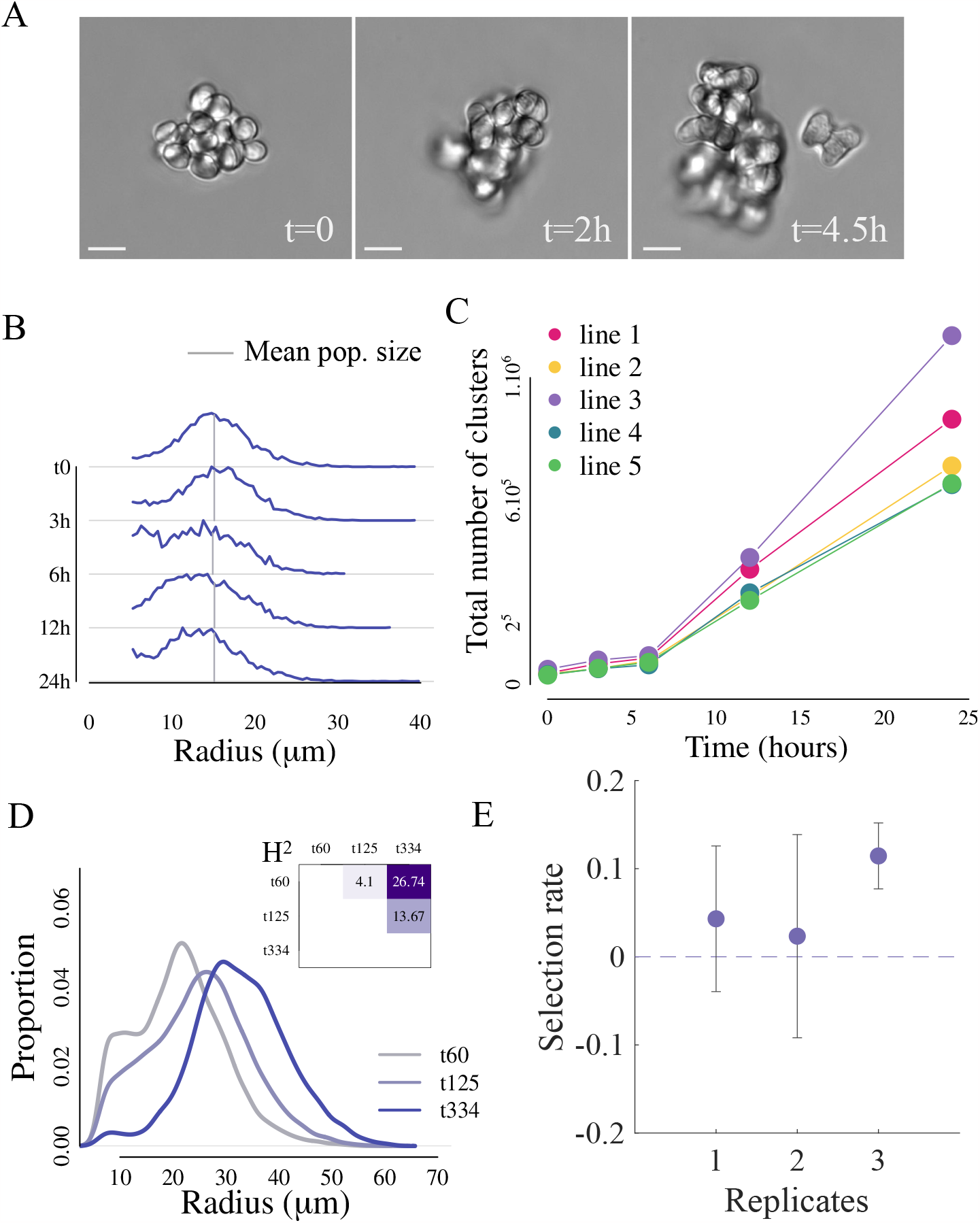
Multicellular *S. pombe* have an emergent multicellular life cycle in which groups become units of selection. (A) *S. pombe* clusters fracture during growth, even without mechanical strain from mixing, producing propagules in soft agar (full time-lapse video in supplementary material, scale bar is 10 *μ*m). (B) The size distribution of the population remains stable over the 24-hour period of growth, during which (C) there is a lag phase followed by a sharp increase in cluster number. (D) Size is heritable: we measured the broad sense heritability (*H*^2^) in all-pairwise comparisons between three time points (t60, t126 and t334, see inset). See Figure A.2 for heritability values for all lines. (E) Group size is a selectable trait: when subjected to settling selection, larger groups (i.e., evolved clusters at t334) do, on average, 6% better than smaller groups (i.e., evolved clusters at t60). All experiments in this figure were conducted using line 3.

Next, to determine if the variation in group size is heritable, we measured the population size of all five populations at three different time points (60, 125 and 334 days, which correspond to ∼140, 300 and 790 generations, respectively) and calculated the broad sense heritability, *H*^2^ (Figure 3 D) for all pairwise differences within lines (t60 vs. t125, t60 vs. t334, t125 vs. t334) using a variance partitioning approach. Broad sense heritability quantifies the proportion of the phenotypic variance in a population that can be attributed to genetic variance. Group size had a heritable component in most pairwise competitions: *H*^2^ ranged from 0.01% to 27% (Figure A.2, 3 D). The highest heritability estimates came from comparisons between t334 strains and their t0 ancestors, and in general *H*^2^ scaled with the amount of evolutionary time separating two isolates.

Finally, to determine whether these heritable differences in cluster size grant a survival benefit during selection for multicellularity, we conducted a competition experiment between two strains with significant differences in size (i.e., 28 *μ*m and 38 *μ*m mean cluster radius for t60 and t334 isolates, respectively, see Figure 2 C). In brief, we measured the frequency of each genotype before and after applying one round of settling selection. As a result, t334 clusters had a 6% higher survival rate than t60 clusters (across four replicate competitions, Figure 3 E). Taken together (Figures 3 A-E), our results demonstrate that evolved *S. pombe* clusters exhibit characteristics of Darwinian individuality: cuboidally packed clusters display a nascent multicellular life cycle that reproduces multicellular propagules (Figures 3 A-C), group size is heritable (Figure 3 D), and these differences in multicellular size confer a survival advantage (Figure 3 E), showcasing the essential components of a major transition in individuality.

To examine the genetic basis of cluster formation in *S*.*pombe*, we sequenced one isolate from each line from the latest time point of our evolution experiment (time 334). Two lines displayed mutations in the *ACE2* gene: a loss of the start codon (line 2) and a frameshift mutation (line 3) (see supplementary Table 2). Mutations in *ACE2* also arose as a mechanism of group formation in response to similar selection in the budding yeast *Saccharomyces cerevisiae*. While both species are yeast, they are distantly related, diverging 330 to 420 million years ago ^36^. This observation exemplifies a case of convergent evolution despite a deep temporal divergence. When we knocked out *ACE2* in the unicellular *S. pombe* ancestor, we observed the emergence of multicellular clusters (Figure 4 A). This mutation alone is not sufficient to fully recapitulate the evolved phenotype: it formed groups with a mean radius that was only 1.4-fold times larger than the unicellular ancestor, which was still 60% smaller than 60-day evolved *S*.*pombe* (mean radius of ancestor cells = 5.9 *μ*m, n = 1750, mean radius *ace2*Δ*::ACE2* = 8.5 *μ*m, mean radius *ace2*Δ*::ace2*Δ= 9.7 *μ*m and n = 1722, mean radius 60-day evolved clusters = 14.1 *μ*m and n = 510, all differences were significant [*<* 0.001] in Tukey’s HSD after one-way ANOVA, *F*_3,8376_ = 1010, *<* 2.10^−16^).

**FIGURE 4.**
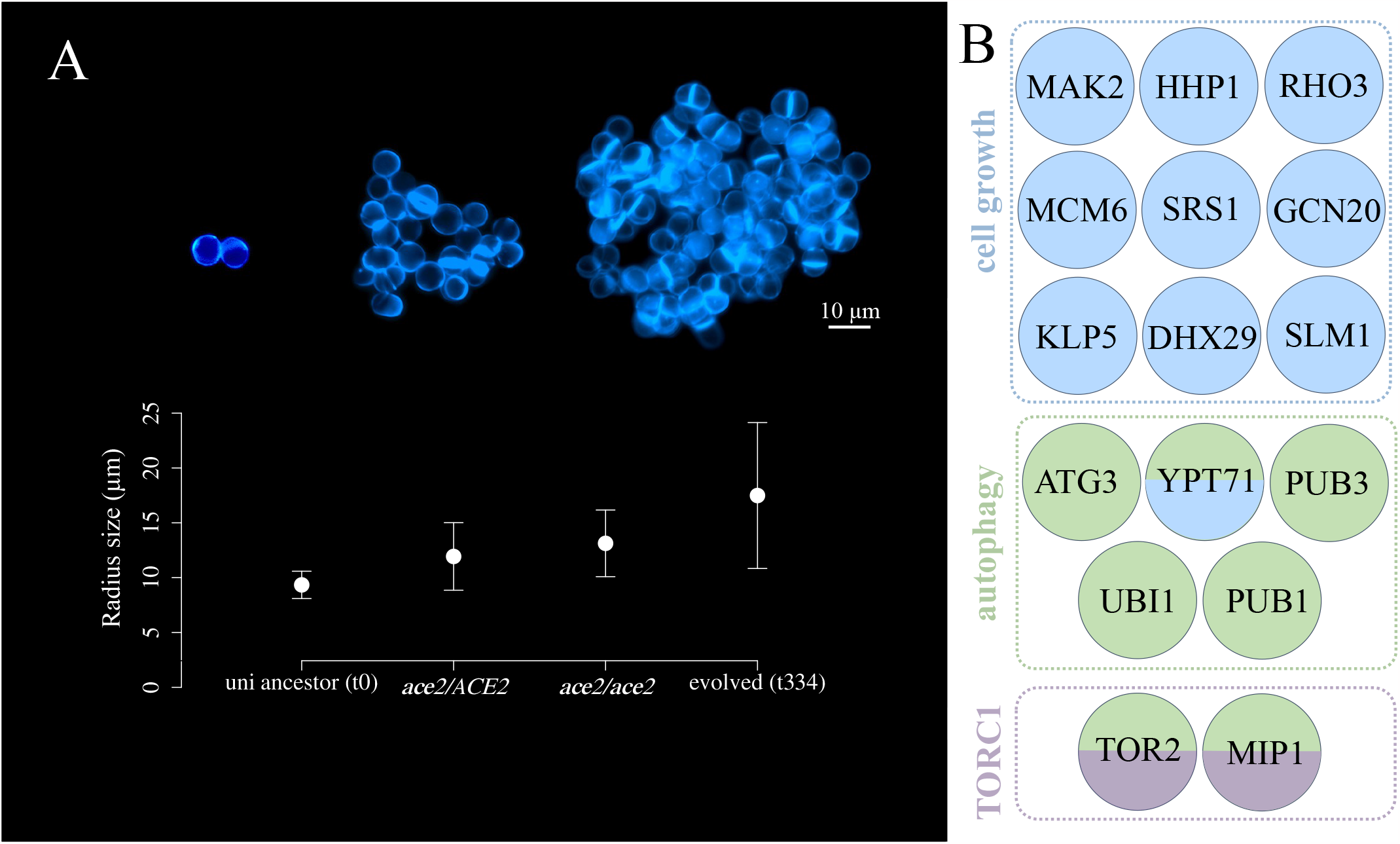
Genetics of multicellular *S. pombe*. (A) We found missense mutations in *ACE2* in two independently evolved lines. Loss-of-function mutations in *ACE2* (both heterozygous and homozygous) resulted in the formation of small multicellular clusters, but did not fully recapitulate the evolved phenotype (scale bar is 10 *μ*m). (B) The GO enrichment analysis identified mutations involved in cell growth, autophagy, and the TORC1 complex. Genes are colored according to their GO category, with two colors indicating genes found in two categories.

GO enrichment analysis on the set of mutations highlighted genes involved in autophagy, cell growth, and in pathways like the target of rapamycin complex 1 (TORC1) (Figure 4 B). For instance, two lines independently evolved mutations in *HHP1* (lines 2 and 4) and in *TOR2* (lines 1 and 3). These genes code for serine/threonine protein kinases that could influence cell growth. Additionally, lines 1 and 5 presented a mutation in *EXG1* gene, a cell wall glucan involved in cell wall organization or biogenesis, which may be involved in cellular adhesion. However, further work will be required to determine what role, if any, these mutations have in the evolution of multicellularity via cuboidal packing in our system.

## 4 DISCUSSION

Cuboidal cell packing is a common yet understudied multicellular morphology that has evolved independently in diverse lineages, from ancient fossils to modern algae, bacteria, and archaea. Previous studies have suggested that cuboidal cell packing may arise from spherical unicellular ancestors that divide by binary fission and alternate their division planes ^14^. However, no prior work has directly examined how this topology arises *de novo*, and whether it can support an evolutionary transition in individuality. Here, we used spherical fission yeast as a model system to experimentally evolve cuboidal cell packing under settling selection, which favors larger size. We chose fission yeast because they are genetically tractable, have a short generation time, and can be easily manipulated to produce spherical cells by mutating genes involved in cell shape determination ^19^.

We observed the emergence of multicellular clusters with cuboidal organization within 20 rounds of selection, displacing the unicellular ancestor. These clusters displayed key hallmarks of an evolutionary transition in individuality: groups possessed an emergent life cycle, their size was heritable, and they responded to group-level selection via multicellular adaptation. The size distribution of the population was remarkably stable, even as the population size increased super-linearly. Combined with our observation of spontaneous fracture during growth (Figure 3 A), this suggests that fracture is driven by cell-cell strain accumulation during growth, as has previously been shown in snowflake yeast ^37^. Like snowflake yeast, this life cycle arises “for free” as an emergent property of cellular crowding during growth.

40% (2/5) replicate populations had loss of function mutations in the gene *ACE2*. This is similar to what was seen with budding yeast, *S. cerevisiae*, when it was evolved under the same selective regime: 50% (5/10) of the replicate lines gained loss-of-function mutations in *ACE2* ^10^. These fungal species last shared a common ancestor 330-420 million years ago ^36^, and have deeply divergent cell biology. This observation exemplifies a case of convergent evolution over at both phenotypic and genotypic levels. Moreover, *ACE2* mutations also induce multicellular group formation in *Candida glabrata* ^38^, another species of budding yeast that is as divergent from *S. cerevisiae* as humans are from fish. *ACE2* plays a conserved role in regulating cytokinesis across the Ascomycota, so perhaps it is not surprising that it would play a similar role in such distantly related organisms.

The convergent evolution of loss of function mutations in *ACE2* highlights the powerful historical contingency exerted by the unicellular ancestor’s cell biology on the mechanisms and dynamics of multicellular evolution. And yet, despite a similar mechanism of group formation (mother-daughter adhesion due to incomplete cytokinesis), budding and fission yeast generate remarkably different multicellular morphologies: budding generates fractal trees, while binary fission generates groups with cuboidal packing. While both topologies generate an emergent multicellular life cycle that allow groups of cells to become Darwinian individuals, little is known about how these early, yet fundamental, differences in group formation will affect their subsequent multicellular evolution.

Some early clues have emerged from ongoing evolution experiments: over 600 transfers with daily settling selection, snowflake yeast evolved to be *>* 20, 000-fold larger, and *>* 10, 000-fold more mechanically tough. They accomplished this by evolving novel biomechanics that leverage their branching phenotype: the evolution of elongate cells reduced the accumulation of cell-cell strain (which when it gets large enough breaks a mother-daughter bond, and breaks the group, ^39^). When combined with mutations strengthening connections between cells, groups withstood fracture long enough that the cells began to grow around one another, ultimately entangling. Through entanglement, which requires many bonds to be broken before a propagule is separated, snowflake yeast leveraged their specific multicellular morphology to drive considerable multicellular trait innovation. It is unclear whether organisms with cuboidal packing can evolve to form large, mechanically tough groups. Consistent with the relatively small size achieved in our experiment here, most extant multicellular organisms with this growth form remain small and simple, though *Archaeophycus* (Figure 1 A, panel 5 is a notable counter-example). Entanglement, which has evolved readily in organisms with a branching morphology, does not appear to be possible in an organism composed of cuboidal subunits. Whether, relative to other multicellular topologies, cuboidal packing faces stronger mechanical constraints on size and toughness, and how this relates to the evolution of multicellularity complexity more broadly, remain open questions ^40^.

## 5 CONCLUSION

In this paper, we explore the evolution of multicellularity through cuboidal packing, a form of clonal multicellularity in which cells stay together after division, forming groups composed of 4 or 8 cell subunits. We show that spherical fission yeast can rapidly evolve multicellular clusters with cuboidal organization under size-based selection, and that these clusters exhibit key features of an evolutionary transition in individuality. We also find that mutations in the *ACE2* gene, which prevents cell separation after division, are responsible for this phenotype in two independent lineages. Remarkably, *ACE2* mutations also underlie the transition to multicellularity in budding yeast, a distantly related lineage. Our findings illustrate the simplicity and convergence of the genetic and biophysical factors that enable the origin of multicellular groups and life cycles, and shed light on the evolution of multicellularity via cuboidal cell packing across the tree of life.

## 7 AUTHOR CONTRIBUTIONS STATEMENT

RMP, GOB and WCR conceived the project. RMP, PCK, DTL, MD, and WW conducted the experiments. RMP analysed the data and generated the figures. RMP, GOB and WCR wrote the manuscript.

## 8 ACKNOWLEDGMENTS

We are grateful to Matthias Sipiczki, for sending us the spherical *S. pombe* ancestor. We would like to thank the Ratcliff lab for thoughtful discussions in the preparation of this manuscript as well as the GT QBioS Graduate Program for its support. This work is supported by the National Science Foundation (Grant No. DEB-1845363).

## 9 SUPPLEMENTARY INFORMATION

Supplementary Tables

All script and raw data will be made available upon publication.

## A SUPPLEMENTARY FIGURES

□

**FIGURE A.1.**
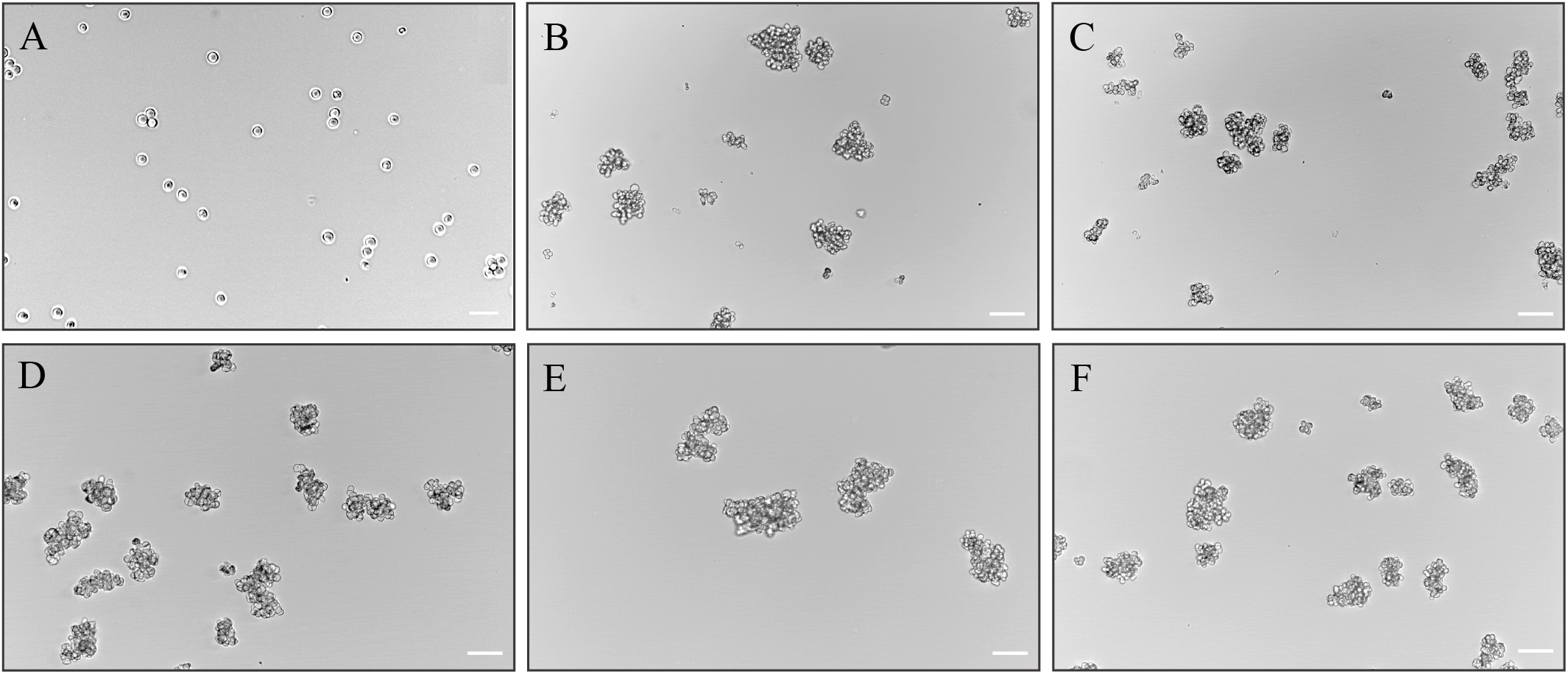
Images showing the ancestor (A) and 334-day evolved populations of *S*.*pombe* (Lines 1-5 corresponding to B-F). Images were taken at a 20X magnification on a Nikon Eclipse Ti-E microscope. Scale bar in (A) is 20 *μ*m. All other scale bars are 50 *μ*m.

**FIGURE A.2.**
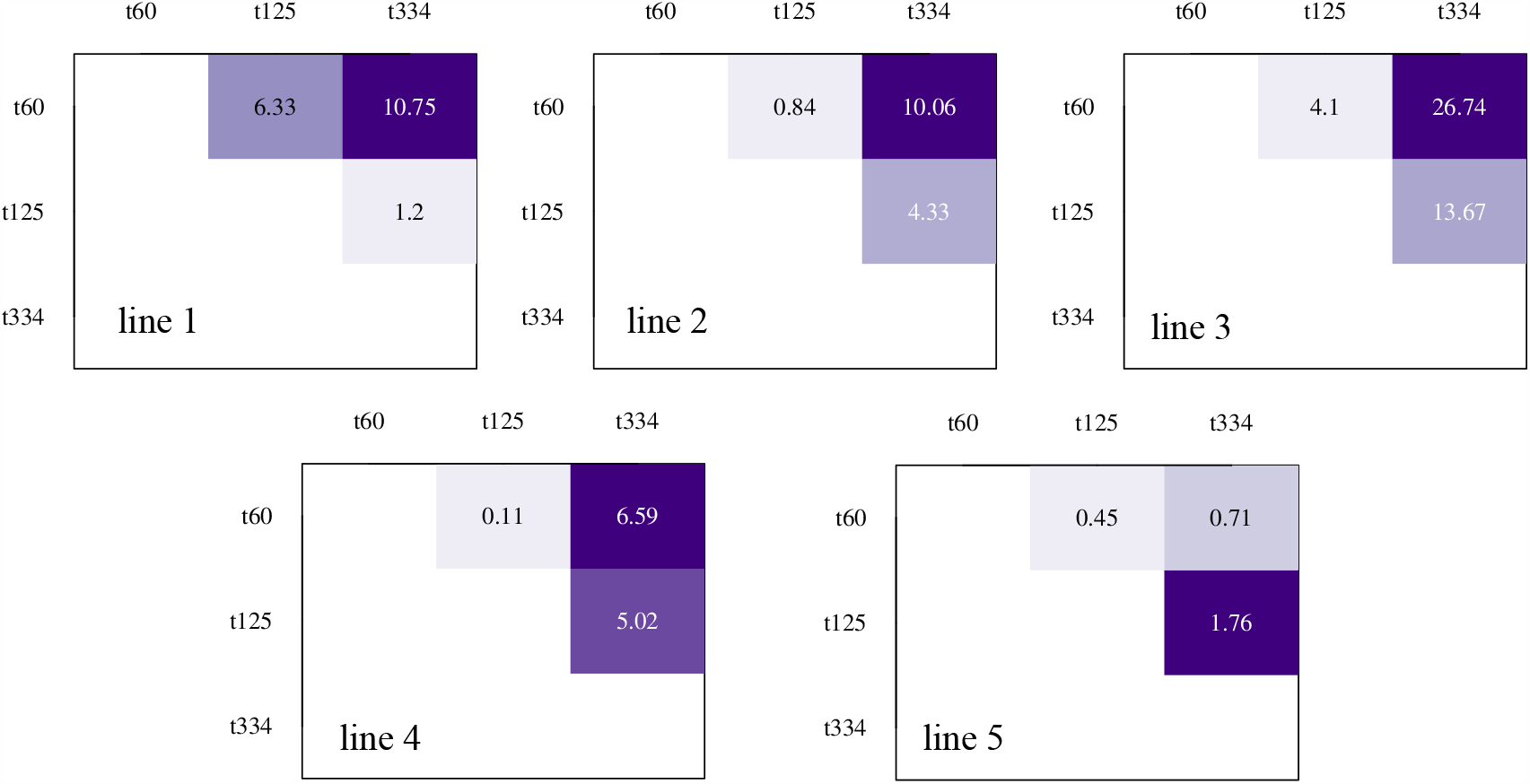
Pairwise broadsense heritability (*H*^2^) values for all pairwise comparisons for t60, t125 and t334 for all 5 independently evolving lines.

**FIGURE A.3.**
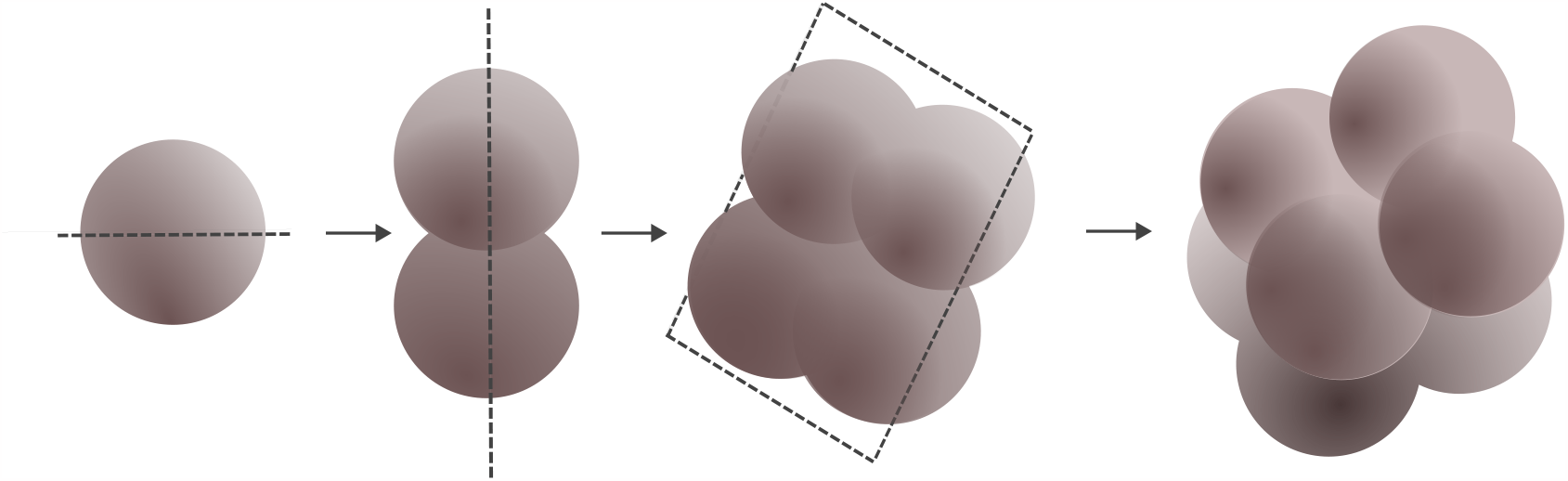
Schematic of the divisions planes leading to cuboidal packing in spherical fission yeast. Each plane of division is perpendicular to the previous plane of division. In the spherical mutant of *S*.*pombe*, the septum does not divide the cell into equally-sized daughter cells, adding stochasticity to this idealized model.

## References

1. Umen J, Herron MD. Green algal models for multicellularity. Annual review of genetics 2021; 55: 603–632.

2. Grosberg RK, Strathmann RR. The evolution of multicellularity: a minor major transition?. Annu. Rev. Ecol. Evol. Syst. 2007; 38: 621–654.

3. Olson BJ, Nedelcu AM. Co-option during the evolution of multicellular and developmental complexity in the volvocine green algae. Current Opinion in Genetics & Development 2016; 39: 107–115.

4. Bernardes JP, John U, Woltermann N, Valiadi M, Hermann RJ, Becks L. The evolution of convex trade-offs enables the transition towards multicellularity. Nature communications 2021; 12(1): 4222.

5. Suga H, Chen Z, De Mendoza A, et al. The Capsaspora genome reveals a complex unicellular prehistory of animals. Nature communications 2013; 4(1): 2325.

6. King N, Westbrook MJ, Young SL, et al. The genome of the choanoflagellate Monosiga brevicollis and the origin of metazoans. Nature 2008; 451(7180): 783–788.

7. Kulkarni P, Behal A, Mohanty A, Salgia R, Nedelcu AM, Uversky VN. Co-opting disorder into order: Intrinsically disordered proteins and the early evolution of complex multicellularity. International Journal of Biological Macromolecules 2022; 201: 29–36.

8. Day TC, Höhn SS, Zamani-Dahaj SA, et al. Cellular organization in lab-evolved and extant multicellular species obeys a maximum entropy law. Elife 2022; 11: e72707.

9. Pentz JT, MacGillivray K, DuBose JG, et al. Evolutionary consequences of nascent multicellular life cycles. eLife 2023; 12: e84336.

10. Ratcliff WC, Fankhauser JD, Rogers DW, Greig D, Travisano M. Origins of multicellular evolvability in snowflake yeast. Nature communications 2015; 6(1): 6102.

11. Mitchison-Field LM, Vargas-Muñiz JM, Stormo BM, et al. Unconventional cell division cycles from marine-derived yeasts. Current Biology 2019; 29(20): 3439–3456.

12. Niklas KJ, Wayne R, Benítez M, Newman SA. Polarity, planes of cell division, and the evolution of plant multicellularity. Protoplasma 2019; 256: 585–599.

13. Xiao S, Muscente A, Chen L, et al. The Weng’an biota and the Ediacaran radiation of multicellular eukaryotes. National Science Review 2014; 1(4): 498–520.

14. Xiao S, Zhang Y, Knoll AH. Three-dimensional preservation of algae and animal embryos in a Neoproterozoic phosphorite. Nature 1998; 391(6667): 553–558.

15. Anderson RP, Macdonald FA, Jones DS, McMahon S, Briggs DE. Doushantuo-type microfossils from latest Ediacaran phosphorites of northern Mongolia. Geology 2017; 45(12): 1079–1082.

16. Proeschold T, Darienko T. The green puzzle Stichococcus (Trebouxiophyceae, Chlorophyta): New generic and species concept among this widely distributed genus. Phytotaxa 2020; 441(2): 113–142.

17. Genome Research oWIC. Methanosarcina acetivorans.

18. DiMaio MA, Park WG, Longacre TA. Gastric Sarcina organisms in a patient with cystic fibrosis. Human Pathology: Case Reports 2014; 1(3): 45–48.

19. Sipiczki M, Yamaguchi M, Grallert A, et al. Role of cell shape in determination of the division plane in Schizosaccharomyces pombe: random orientation of septa in spherical cells. Journal of bacteriology 2000; 182(6): 1693–1701.

20. Murray JM, Watson AT, Carr AM. Transformation of Schizosaccharomyces pombe: Lithium Acetate/Dimethyl Sulfoxide Procedure. Cold Spring Harbor Protocols 2016; 2016(4): pdb–prot090969.

21. Chen S, Zhou Y, Chen Y, Gu J. fastp: an ultra-fast all-in-one FASTQ preprocessor. Bioinformatics 2018; 34(17): i884–i890.

22. Li H. Aligning sequence reads, clone sequences and assembly contigs with BWA-MEM. arXiv preprint arXiv:1303.3997 2013.

23. Li H, Handsaker B, Wysoker A, et al. The sequence alignment/map format and SAMtools. bioinformatics 2009; 25(16): 2078–2079.

24. Barnett DW, Garrison EK, Quinlan AR, Strömberg MP, Marth GT. BamTools: a C++ API and toolkit for analyzing and managing BAM files. Bioinformatics 2011; 27(12): 1691–1692.

25. Auwera V. dGA, Carneiro MO, Hartl C, et al. From FastQ data to high-confidence variant calls: the genome analysis toolkit best practices pipeline. Current protocols in bioinformatics 2013; 43(1): 11–10.

26. Danecek P, Auton A, Abecasis G, et al. The variant call format and VCFtools. Bioinformatics 2011; 27(15): 2156–2158.

27. Robinson JT, Thorvaldsdóttir H, Turner D, Mesirov JP. igv. js: an embeddable JavaScript implementation of the Integrative Genomics Viewer (IGV). Bioinformatics 2023; 39(1): btac830.

28. García P, Tajadura V, García I, Sánchez Y. Rgf1p is a specific Rho1-GEF that coordinates cell polarization with cell wall biogenesis in fission yeast. Molecular biology of the cell 2006; 17(4): 1620–1631.

29. Ratcliff WC, Denison RF, Borrello M, Travisano M. Experimental evolution of multicellularity. Proceedings of the National Academy of Sciences 2012; 109(5): 1595–1600.

30. Dudin O, Wielgoss S, New AM, Ruiz-Trillo I. Regulation of sedimentation rate shapes the evolution of multicellularity in a close unicellular relative of animals. PLoS biology 2022; 20(3): e3001551.

31. Ratcliff WC, Herron MD, Howell K, Pentz JT, Rosenzweig F, Travisano M. Experimental evolution of an alternating uni-and multicellular life cycle in Chlamydomonas reinhardtii. Nature communications 2013; 4(1): 2742.

32. Driscoll WW, Travisano M. Synergistic cooperation promotes multicellular performance and unicellular free-rider persistence. Nature communications 2017; 8(1): 15707.

33. Godfray HCJ, May RM. Open questions: are the dynamics of ecological communities predictable?. BMC biology 2014; 12(1): 1–3.

34. Godfrey-Smith P, Bouchard F, Huneman P. Darwinian individuals. From groups to individuals: Evolution and emerging individuality 2013; 16: 17.

35. Rose CJ, Hammerschmidt K. What do we mean by multicellularity? The evolutionary transitions framework provides answers. Frontiers in Ecology and Evolution 2021; 9: 730714.

36. Sipiczki M. Where does fission yeast sit on the tree of life?. Genome biology 2000; 1(2): 1–4.

37. Jacobeen S, Pentz JT, Graba EC, Brandys CG, Ratcliff WC, Yunker PJ. Cellular packing, mechanical stress and the evolution of multicellularity. Nature physics 2018; 14(3): 286–290.

38. Kamran M, Calcagno AM, Findon H, et al. Inactivation of transcription factor gene ACE2 in the fungal pathogen Candida glabrata results in hypervirulence. Eukaryotic cell 2004; 3(2): 546–552.

39. Bozdag GO, Zamani-Dahaj SA, Day TC, et al. De novo evolution of macroscopic multicellularity. Nature 2023: 1–8.

40. Knoll AH. The multiple origins of complex multicellularity. Annual Review of Earth and Planetary Sciences 2011; 39: 217–239.

